# Bioelectronic microfluidic wound healing

**DOI:** 10.1101/2022.07.08.499276

**Authors:** Sebastian Shaner, Anna Savelyeva, Anja Kvartuh, Nicole Jedrusik, Lukas Matter, José Leal, Maria Asplund

**Affiliations:** Department of Microsystems Engineering, University of Freiburg, Freiburg, Germany; Brainlinks-Braintools Center, University of Freiburg, Freiburg, Germany; Freiburg Institute for Advanced Studies (FRIAS), University of Freiburg, Freiburg, Germany; Division of Nursing and Medical Technology, Luleå University of Technology, Luleå, Sweden; Department of Microtechnology and Nanoscience, Chalmers University of Technology, Gothenburg, Sweden

**Keywords:** electrotaxis, galvanotaxis, direct current electric field, wound healing, microfluidics, conducting polymers

## Abstract

This work delves into the impact of direct current (DC) stimulation on both healthy and diabetic *in vitro* wound healing models of keratinocytes, the most prevalent cell type of the skin. The augmentation of non-metal electrode materials and prudent microfluidic design allowed for a platform to study the effects of different sustained (12 hours DC) electric field configurations on wound closure dynamics. We found that electric guidance cues (≃ 200mVmm^−1^) enhance wound closure rate by nearly 3X for both healthy and diabetic-like keratinocyte sheets, compared to their respective controls. The motility-inhibited keratinocytes regained wound closure rates with stimulation (increase from 1.0 to 2.8% hr^−1^) comparable to healthy non-stimulated keratinocyte collectives (3.5% hr^−1^). Our results bring hope that electrical stimulation is a viable pathway to accelerate wound repair.

## Introduction

For most of us, a wound is a minor nuisance, which heals itself without much to any conscious effort. However, for people with certain chronic diseases (e.g, diabetes mellitus, peripheral vascular disease), compromised immune systems (e.g., systemic lupus erythematosus), or even with common systemic factors, such as poor nutrition and aging, acute wounds are more prone to become chronic. In fact, the high prevalence of chronic wounds constitute an enormous socioeconomic burden (≈ 1 to 3 % of total healthcare spending in developed countries and growing as the median age of populations grow older)^1^, as well as suffering for the actual patients.^2, 3^ Strategies to promote faster healing for these patient groups are therefore urgently needed. The healing process is often categorized in four sequential, yet overlapping phases: hemostasis, inflammation, growth, and maturation. There are many cell types involved in these concurrent phases (in general order of appearance): activated platelets, neutrophils, monocytes, macrophages, mast cells, dendritic cells, T cells, endothelial cells, pericytes, hematopoietic stem cells, fibroblasts, myofibroblasts, melanocytes, and keratinocytes.^4^ In the wound healing process these cell types are recruited, some already being residents of the wound site while others have to traverse long distances via the circulatory system.

Chemical, mechanical, and electrical gradients all contribute to recruiting or guiding the aforementioned cells to the wound: processes called chemotaxis,^5^ haptotaxis/durotaxis,^6, 7^ and electrotaxis/galvanotaxis,^8^ respectively. Notably, electrotaxis refers to the ability of cells to align their migration with electric fields (EFs). Neutrophils^9^, monocytes^10^, lymphocytes^10^, macrophages^11^, endothelial cells^12^, fibroblasts^13^, and keratinocytes^14^ have all been revealed to be electrotactic. Interestingly, wounds naturally form small EFs when the skin’s epithelial layer is broken. This transepithelial potential (~ 10 to 60 mV), which actively pumps sodium ions (Na^+^) basally inwards and chlorine ions (Cl^−^) apically outwards, is short-circuited after injury where positive current flows radially towards the wound center.^15^ There is large inter-individual variability in the strength of this naturally-occurring (i.e., endogenous) EF, which depends on the systemic nature of the patient (e.g., age, disease). For example, it was shown that the lateral EF of 18-25-year-olds (107 to 148 mVmm^−1^) is nearly 48% larger compared to 65-80-year-olds (56 to 76 mVmm^−1^).^16^ Taken together with the fact that most skin cells exhibit electrotactic ability, the discovery of these endogenous EFs at wounds have led to the hypothesis that electrical cues are important for migratory processes in wound healing.^17, 18^ This hypothetical precedent still remains an ever-evolving topic of interest, especially since earlier work has not fully unraveled the merit of EF magnitude and distribution on electrically-guided closure of actual wounds.

The most prevalent cell type in the skin, keratinocytes, are densely packed within any given lateral layer, and are also tightly arranged in vertical tiers (i.e., stratified) where they become more differentiated the closer they get to the outermost, apical layer. In the skin, as well as in confluent cultured cell layers, keratinocytes migrate as a collective.^19^ Prior *in vitro* studies on electrotaxis of skin cells have typically focused on single cells, thus neglecting how the complex organization of cells in actual skin impact the migratory behavior. Whereas, collective cell migration is more indicative of in vivo cell dynamics for cell types like keratinocytes. On a group level, coordination within the cell collective starts with the protrusion of cells at the group’s edge (i.e., leader cells) and is propagated through cell-substrate forces (i.e., traction of cell membrane-bound focal adhesions with the substrate)^20, 21^ and cell-cell forces (i.e., normal and shear forces via adherens junctions).^22, 23^ It is only recently that there has been more focus on electrotaxis-mediated group migration.^24, 25^ For instance, Zajdel *et al*.^26^ showed that two patterned monolayers of keratinocytes, with a 1.5 mm gap, can be influenced by an EF stimulus (i.e., 200 mV mm^−1^) and close the gap between the two epithelial sheets twice as fast as compared to non-stimulated controls. This work set an important precedent, and calls for further translational efforts to make EF-accelerated repair accessible to patients who are at risk for chronic wounds. A crucial step in this process would be to more closely replicate the process of wounding, where mechanical stress (e.g., physical removal of cells) induces ATP and gap junction-mediated calcium waves across the monolayer of keratinocytes.^27^. Furthermore, as healthy skin typically heals well, it is paramount to analyse this effect in cultured disease models, which are associated with impaired wound healing and keratinocyte motililty.

In this work, the aim was threefold: (a) explore the influence of electrical guidance cues (EF distributon and magnitude) on the rate of wound closure, (b) establish a new DC stimulation electrode material that circumvents common pitfalls of standard materials, (c) and establish a diabetic wound model to examine if electrical stimulation improves otherwise poor wound closure dynamics. In order to facilitate this, we developed a microfluidic version of the ‘classical’ scratch wound assay, which allows us to explore the parameter space in which EF stimulation accelerates wound repair. Multiple fluidic concepts are analyzed to identify the layout that best mimics the standard scratch assay method, but with the superior experimental control provided by the microfluidic platform. Direct current (DC) compatible electrode materials are a key ingredient for EF stimulation *in vitro*, and will furthermore be essential for clinical translation.^28, 29^ Here we show that electrodes based on a combination of laser-induced graphene (LIG) and PEDOT:PSS hydrogel, integrated within the platform, were capable of sustaining DC stimulation over hours. This is not only important for the *in vitro* application, but likewise a prerequisite for subsequent clinical translation of the concept. Leveraging this wound-on-a-chip environment, we were able to explore the electrical wound healing concept, first for healthy cells and then using a culture-based model mimicking diabetes. Not only were we able to demonstrate that EF stimulation was effective for accelerating the wounds for the healthy cells but, importantly, could restore the impaired mobility of diabetic-like cells as well.

## Results

### Microfluidic design for tailoring electric field distributions around scratch

The protocol for making the electrotaxis scratch device leverages using a common prototyping equipment (i.e., CO_2_ laser) for both microfluidic adhesive structuring^30^ and electrode fabrication^31^ (Fig. 1 a,b, see Methods section for more details). The central challenge of developing a scratch assay protocol inside a microfluidic channel is how to seed, grow, and then scratch a monolayer of cells all within an enclosed domain. Non-contact techniques of creating a scratch in a closed channel, such as UV exposure through a shadow mask or using a laser to ablate cells, fail to result in reproducible conditions as unsuccessful removal of debris have propagating negative effects on the leading edges of the scratch. For this reason, we developed an approach that allowed for cells to be seeded onto an adhesive-masked Petri dish where the cells adhere to the exposed area of the dish. Using a double-sided adhesive with protective liners on both sides allows for removal of the bottom-side liner to adhere the exposed adhesive to the dish all the while keeping on the top liner to protect the top adhesive from getting wet, thus acting as a sacrificial layer. After seeding and overnight incubation (Fig. 1b step 3, Supplemental Fig. 5), monolayers are injured in the traditional way (i.e., mechanical removal via pipette tip).^32^ We found that wedging a vacuum aspirator metal tip into a sterile plastic pipette tip allowed for more reproducible removal of cells with less cellular debris left in the wake of the scratch path. After scratching, the adhesive’s top liner is removed so that a plastic lid with reservoirs could be manually aligned and pressed onto the exposed adhesive to complete the enclosure of the microchannels, then immediately filled with media (Fig. 1b steps 2 & 6). The reservoirs are only connected fludically via the microchannels. A set of non-metal conducting hydrogel electrodes (i.e., PEDOT:PSS hydrogel-coated laser-induced graphene) are assembled with custom 3D-printed pogo pin electrical connection adapters and holders. Note that the circles at the end of each branching microchannel act as the beginning of the open reservoirs, such that only the four branches are confined to the micron-size geometry in all dimensions.

**Figure 1.**
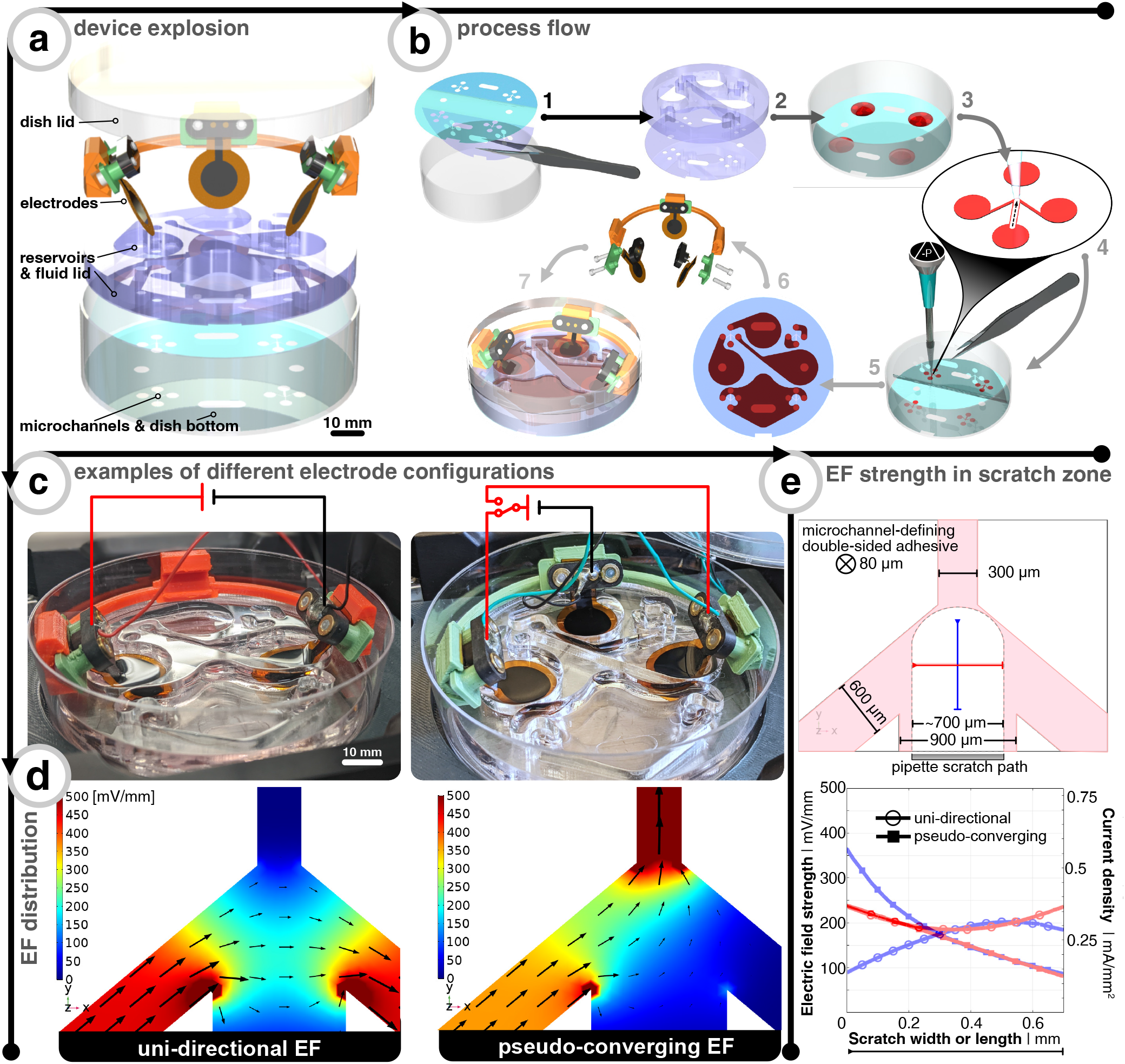
Microfluidic design to allow different electric field (EF) distributions around scratch. (a) explosion view of device showing the main components. (b) fabrication process flow where the steps are as follows: 1 = bottom liner is peeled off of the laser-cut double-sided adhesive so it can be placed onto a polystyrene dish. 2 = two-part laser-cut acrylic lid is solvent bonded together. 3 = keratinocytes are seeded at high density over open microchannels for 4 hours, then filled with media overnight. 4 = cells are scratched away using a pipette tip fitted around a vacuum aspirator. 5 = top liner is peeled off of the adhesive. 6 = lid from step 2 is aligned and bonded, then channels and reservoirs are filled with cell culture media. 7 = electrodes are assembled with pogo pin adapters and 3D-printed holders. 8 = electrodes are placed into reservoirs and covered with dish lid. (c) Assembled electrotaxis device showing two different electrode configurations where the red (anode) and black (cathode) traces show the electrical connection layout connected to constant current source. The pseudo-converging EF case makes use of a relay to alternate anodes, thus a mirror EF distribution along the y-axis (d) Finite element analysis of EF distribution within microchannel for both electrode configurations where black arrows show the current direction and magnitude. (e) Top illustration shows the dimensions of the microchannel and the typical profile of scratch. The blue and red lines are exemplary profiles to note the EF strength at the center of the scratch in x and y. The bottom graph shows the aforementioned blue and red profiles for both electrode configurations. Note that an electrode input current of 25 and 20 μA is chosen so that center of the channel is about 200mVmm^−1^ for uni-directional and converging cases, respectively.

Wounds found *in vivo* are such that the effective EF points radially towards the wound center.^33^ Electrically-speaking, the wound center acts like a current sink (i.e., cathode) surrounded by an ionic current source (i.e., anode).^15^ When designing the microfluidic device, it is this principle that we mimic. An ideal setup would involve having an infinitesimally-small cathode at the wound center such that all EF vectors guide electrotactic cells to the center. However, the limited ionic charge storage capacity of the electrode materials prevent such miniaturization. An alternative to miniaturizing the cathode, which can visually occlude the closing wound, is to explore how the layout of the microfluidic system can be tailored to allow EF stimulation to converge towards the wound center. Instead, our microfluidic design included a merging microfluidic network where the scratched cells will be centered and different combinations of electrode configurations around scratch can be explored.

The microfluidic network is shaped in a “peace sign” configuration where there are four branches of different widths that lead to individual reservoirs (Fig. 1d). The angled position of the left and right branches help steer current away from an EF “dead zone” at the center of the wound, which would be seen if these branches are perpendicular to the top branch (Supplemental Fig. 1). The purpose of the lower branch is to act as a guide for scratching the cells, where the width (900 um) is chosen so that a p10 pipette tip (≃ ∅ 700 um) could be used as the scratching tool, which is a typical way^32^ to create *in vitro* wounds (Fig. 1e). Secondarily, this lower branch yields the option to have perfusive hydrostatic flow in order to ensure culture health throughout days-long experiments. In order to demonstrate this additional functionality, we used different colored dyes to validate that an effective converging flow can be accomplished without the use of any active component (i.e., no perfusion pump or similar needed) (Supplemental Fig. 2). However, for all functional testing, the perfusive flow option was not utilized as to decouple cell migration due to replenishing nutrients versus applied EF. Two configurations were analyzed in this study. The first will be referred to as the “uni-directional EF” and correspond to an EF across the wound (i.e. not a converging field). This strategy is the most straight forward to implement, as it requires only two electrodes, which are placed at either side of the wound (Fig. 1c,d left column). The electrotactic effect in this case will mainly apply to one side of the wound. According to the electrotactic behavior seen in single cell cultures, the other side would migrate away from the scratch, if only electrotaxis-related forces would dominate the migratory behaviour. However, the majority of studies that explore electrical wound healing *in vivo* exploit this type of setup, despite that it does not fully account for electrotaxis to act on both edges of the wound.^34, 35^ The second configuration is here referred to as the “pseudo-converging EF” profile since there is an anode on each side of the scratch where a timed relay can switch between the two anodes to push the scratch close from both sides in an alternating fashion (Fig. 1c,d right column). Consequently, in this scheme, the electrotactic effect will apply symmetrically to both sides of the scratch. Additionally, the disconnected and polarized anode can passively recharge with ions from solution while the other anode is delivering charge.^29^

Using finite element analysis (FEA) of the three-dimensional microfluidic device, we identified the input current needed to achieve ≃ 200mVmm^−1^ at the center of the scratch, which has been shown to be an optimal EF strength for *in vitro* keratinocyte electrotaxis.^36–38^ As validated by the FEA model, the geometric confinement around the monolayer of scratched cells provided by the microchannel allows for precise control of the field distribution (Fig. 1d). The current and normalized EF direction is indicated via the black arrows and the EF magnitude is depicted via the scaled color gradient and size of the black arrows. In order to visualize the EF magnitude along and across the scratch, two profile lines are drawn (blue and red, respectively in Fig.1e). The plot shows EF magnitude and corresponding microchannel current density along these lines for a given electrode input current (25 μA and 20 μA for uni-directional and pseudo-converging cases, respectively). The hollow circles show that the uni-directional case yields a more uniform EF distribution across (i.e., x-direction, red) and along (i.e. y-direction, blue) compared to the pseudo-converging case. The center of the scratch is defined to be where both lines intersect (for a comparison to non-angled branched designs, see Supplemental Fig. 1). The input currents were selected to match a value of 200mVmm^−1^ at this intersection point.

### Minimal pH shifts and joule heating during direct current stimulation

For reproducibly generating electrotactic behavior, it is fundamental that electrotactic effects are decoupled from other possible interferences. For instance, faradaic reactions at the electrode-electrolyte interface can induce redox reactions, which yield a lowering of pH at the anode (i.e., more H^+^) and raising of pH at the cathode (i.e., less H^+^). Since previous studies have explicitly pointed to pH as a determinant factor of electrotaxis^39^ and since the confined volume inside the microfluidic compartment yields the system more sensitive to such variances, we explicitly validated pH stability within the microfluidic system under DC stimulation. In order to monitor pH, the microchannel floor is coated with the colorimetric pH-sensitive polymer polyaniline (i.e., PANI) (Fig. 2, Supplemental Video 1). The color changes with the oxidation state of the conjugated polymer. At low pH (≃ 3 to 5), the PANI is in its emeraldine salt oxidation state and gives a gradient color from yellow-green to dark green as the pH increases. At more neutral pH (≃ 6 to 8), the PANI gradually changes to the emeraldine base oxidation state and begins to transition from dark green to green-blue to blue. Finally, at higher pH (≃ 9 to 12), the PANI progresses to a fully oxidized state called pernigraniline, where the color goes from blue to dark blue to dark purple.^40^

**Figure 2.**
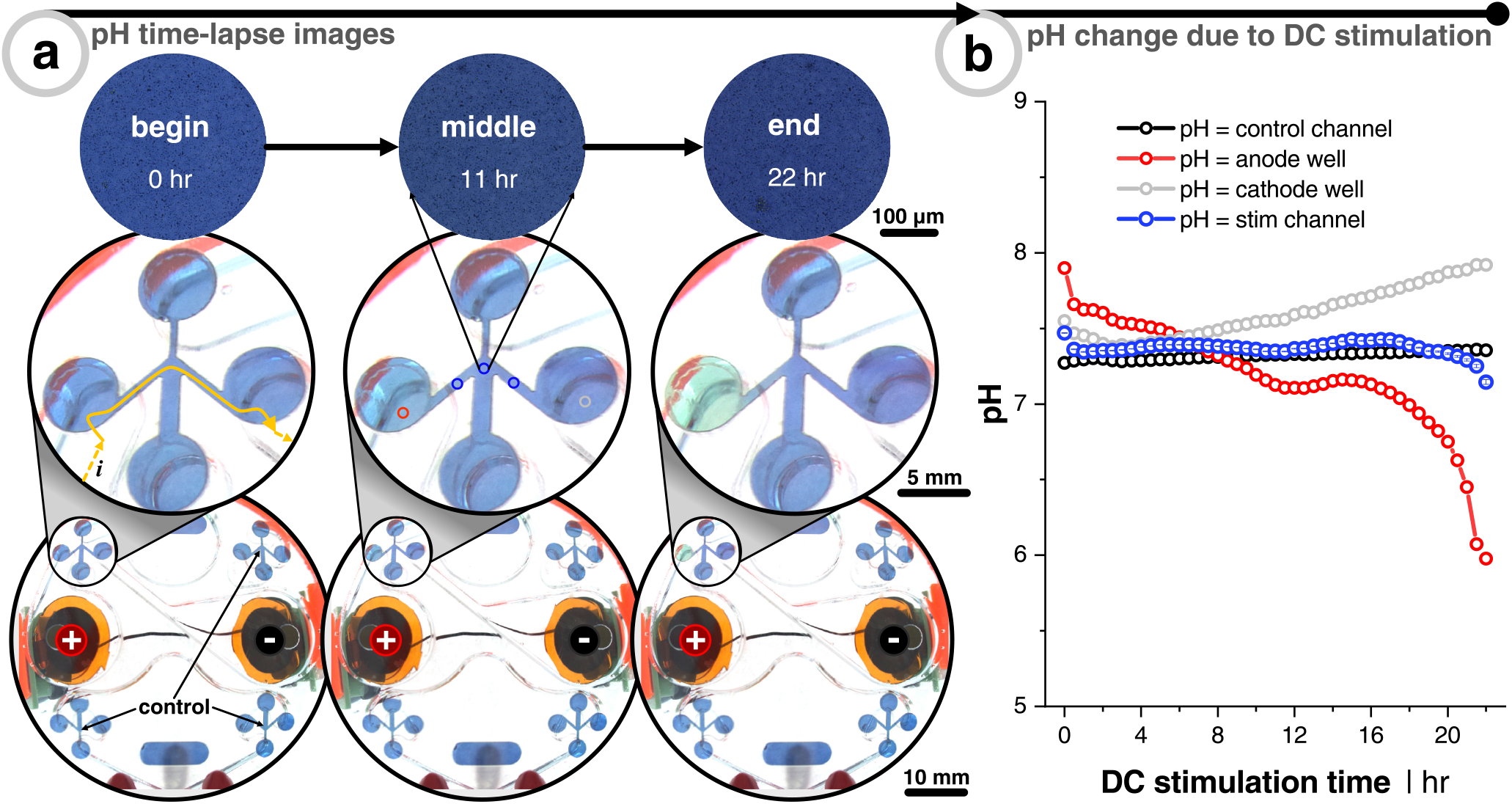
Measuring pH shifts due to direct current (DC) stimulation. (a) Time-lapse example images of 1X phosphate-buffered saline (PBS) filled device with polyaniline (PANI) coated microchannels. The stimulation protocol is a uni-directional EF configuration and stimulated for 22 hr at 25 μA (see Fig. 1). Yellow arrows show current path from reservoir down into and across the microchannel. Note that the images on the bottom two rows are taken with a wide-field camera, whereas the top row is taken with a 5X objective in an incubated (37 °C & 5 % CO_2_) microscope. All quantitative data used are of the 5X images to minimize input light fluctuations. (b) Quantitative output of pH changes as function of DC stimulation time. Colors correspond to the imaging locations shown in (a) of the center image. Black trace is non-stimulated control microchannel. Blue trace is the average of 3 locations (before, center, and after scratch zone) of stimulated microchannel. Red and gray traces are at the interface between reservoir and microchannel entry/exit, which show the anode and cathode wells, respectively. Note that the relationship between color (i.e., hue) and pH is done via a calibration curve of PANI-coated dishes exposed to 15 known pH buffers (Supplemental Fig. 3a,b).

The purpose of the PANI coating is to identify at which cutoff time the DC stimulation will induce notable pH shifts in the microchannel for the given input currents. First, PANI’s pH-sensitivty was tested by coating a thin layer onto small Petri dishes and filling them with 15 known pH buffers to create a calibration curve relationship between the pH and the consequent PANI color (i.e., hue) (Supplemental Fig. 3a,b). The sensitivity to indicate pH change during 20 s of DC stimulation was first verified using a non-buffered saline solution (0.9 %) with relatively high current density (5.7 mAcm^−2^), which is about 250X greater than what was planned for the microchannels) in order to rapidly see pH changes below the electrode pair. As soon as 10 s into stimulation, the area under the cathode and anode begins to turn more basic (purple) and acidic (green), respectively (Supplemental Fig. 3c). This demonstrated PANI’s ability to display the rapid visual pH dynamics as a function of electrochemical faradaic by-products before moving into the device architecture.

Phosphate-buffered saline (PBS) solution of pH 7.4 is used as the testing electrolyte, which also has a similar osmolarity to the keratinocyte media. The PANI is coated in the microchannel analogous to the way cells are seeded in the device (Fig. 1b, step 3 & Fig. 2). We here focused on the uni-directional EF case, which represents the highest current injected into the system, and therefore can be expected to correlate with stronger potential pH shifts. Also, the pseudo-converging EF does not induce an acidic pH swing to the same degree due to minimizing the injected faradaic current by way of using two anodes that rely on relay-switching and passive ion recharging (Supplemental Fig. 3e). Imaging is performed in two ways. The first experiment was with a transmission light microscope fitted with a 5X objective to control light intensity and minimize ambient fluctuations (Fig. 2a, top row). Second, with a wide angle camera lens to concurrently capture a global view of all control and stimulation channels (Fig. 2a, middle and bottom rows). As expected, the PANI at the transition between the anode reservoir and microchannel (i.e., left circle & channel) turns more acidic and vice versa for the cathode well (Supplemental Video 2). From this we conclude that at least 12 hr of DC stimulation is possible without inducing significant pH shifts for this combination of input current, electrode size, stimulation configuration, pH-buffering capacity, and microfluidic design. We would here like to emphasize that no cross-flow was used to dilute potential electrochemical by-products, and also no salt-bridges nor any other supporting system, which otherwise are essential components when using other metal-based electrode systems.

Another possible interference of applying DC across a fluidic resistor (i.e., microchannel) is that joule heating effects could increase metabolism of the cells and allow them to migrate faster. In order to account for this, it is important to have an idea of the amount of joule heating as a function of DC stimulation time. Thermocouples are susceptible to the effects of electromagnetic fields, particularly induced voltages from the EF, and are typically physically much larger than the microchannels employed in this work. Therefore, we opted to focus on a FEA simulation-based approach. Specifically, joule heating effects within the electrolyte were computed using the multiphysics coupling of electromagnetic time-independent equations (current conservation based on Ohm’s law and scalar electric potential) and heat transfer time-dependent equations (energy conversation using Fourier’s law). Both uni-directional and converging cases are explored within a 12 hr stimulation cutoff. The model involved simplifying the geometry to just the stimulation microchannel network, connected reservoirs, surrounding plastic substrate and lid, as well as the electrodes sitting on top of the reservoirs (Supplemental Fig. 4). Even after an energy transfer of 2.74 J (25 μA, 12 hr, 101.6 kΩ) and 1.87 J (20 μA, 12 hr, 108.3 kΩ) for uni-directional and converging EF cases, respectively, the temperature in the scratch zone only rises less than 0.1 °C. Since more current is forced through the smallest width microchannel branch in the pseudo-converging EF case, the electrical resistance was higher leading to a higher joule heating (maximum temperature increase was 0.03 °C and 0.11 °C after 12 hours of DC stimulation for uni-directional and pseudo-converging cases, respectively). Using the combined experimental and computational approach to account for pH shifts and joule heating due to DC stimulation, we could conclude that the cells are safe from electrochemically-induced faradaic reactions and heat-induced apoptosis during the 12 hr stimulation of cells in our platform at these currents.

### DC stimulation expedites wound closure of keratinocytes

Preliminary electrotaxis studies of single cell keratinocytes are in consensus that there is cathodic directionality of migration, but there are mixed reports on small or significant increases of migration speed compared to non-stimulated controls.^37, 38, 41^ In the body, however, keratinocytes are organized as packed layers, and only few studies have attempted to study the more skin-relevant situation of electrotaxis in confluent cell layers. Recently, it was shown that monolayers of keratinocytes have both cathodic migration directionality and increased migratory speed (~ 3X) when subjected to an external direct current EF (200mVmm^−1^).^36^ Globally, collective cells move like an elastic material with a constant tug-of-war between the cells at the advancing edge (i.e., leader cells) and the conglomerate of cells behind them (i.e., follower cells), where there is an interplay of forces that act on the individual and collective group levels.^23^ This is where collective and individual cell migration differ as the latter is not directly influenced by their neighbors, thus highlighting the importance of studying the more wound-realistic crowd migration of damaged epithelial sheets whose leader cells are steering the way.

Healthy keratinocytes are seeded, grown to full confluency, scratched, and DC stimulated according to Fig. 1. The stimulation protocol was either 12 hr of uni-directional or pseudo-converging EF, and each stimulation replicate had multiple internal non-stimulation controls (Fig. 3). Remarkably, DC stimulation resulted in faster scratch closure in all cases. At the end of stimulation, the scratches subjected to uni-directional EF (n = 3, in blue) were ≈ 100% closed, converging EF (n = 3, in red) was ≈ 72% closed, and the controls (n = 9, in black) ≈ 42% closed. For the uni-directional EF case, the effect was even stronger and the full scratch closure was seen even at 10 hr when controls were only ≈ 36% closed, nearly a 3X increase in closure rate. If the effect was purely following the logic of electrotactic behaviour in single cells, one would expect that the closure speed would be faster for the converging EFs, where both edges of the scratch experience a field which should drive them towards the wound center. The faster closure speeds was here instead seen for the uni-directional EF. We hypothesize that this is due to time-varying shifts in the polarization direction of the leader cells, which could have compounding negative traction effects each time the anode is switched. This compounding effect could also explain why the initial slopes of closure rates were similar for both stimulation schemes, but ultimately slows down for the pseudo-converging EF case. Note that the pseudo-converging EF stimulation scheme involved a relay-based switching of the left and right anode every 30 min to allow for cyclic passive recharging of the non-connected and depleted anode with cations from solution, which can be seen in the rapid rising potential at the beginning of each discharge that is predominately due to capacitive current (Fig. 3c). This scheme allowed the electrodes to operate at a lower potential, which would be beneficial to reduce electrochemical side-effects. There is a subtle increase of the potential over time, which is likely due to the sudden switching of anodes adding stress on the cathode. Owing to the superior wound closure performance of the uni-directional EF scheme, it was opted to be the focal stimulation case for diabetic model of keratinocytes.

**Figure 3.**
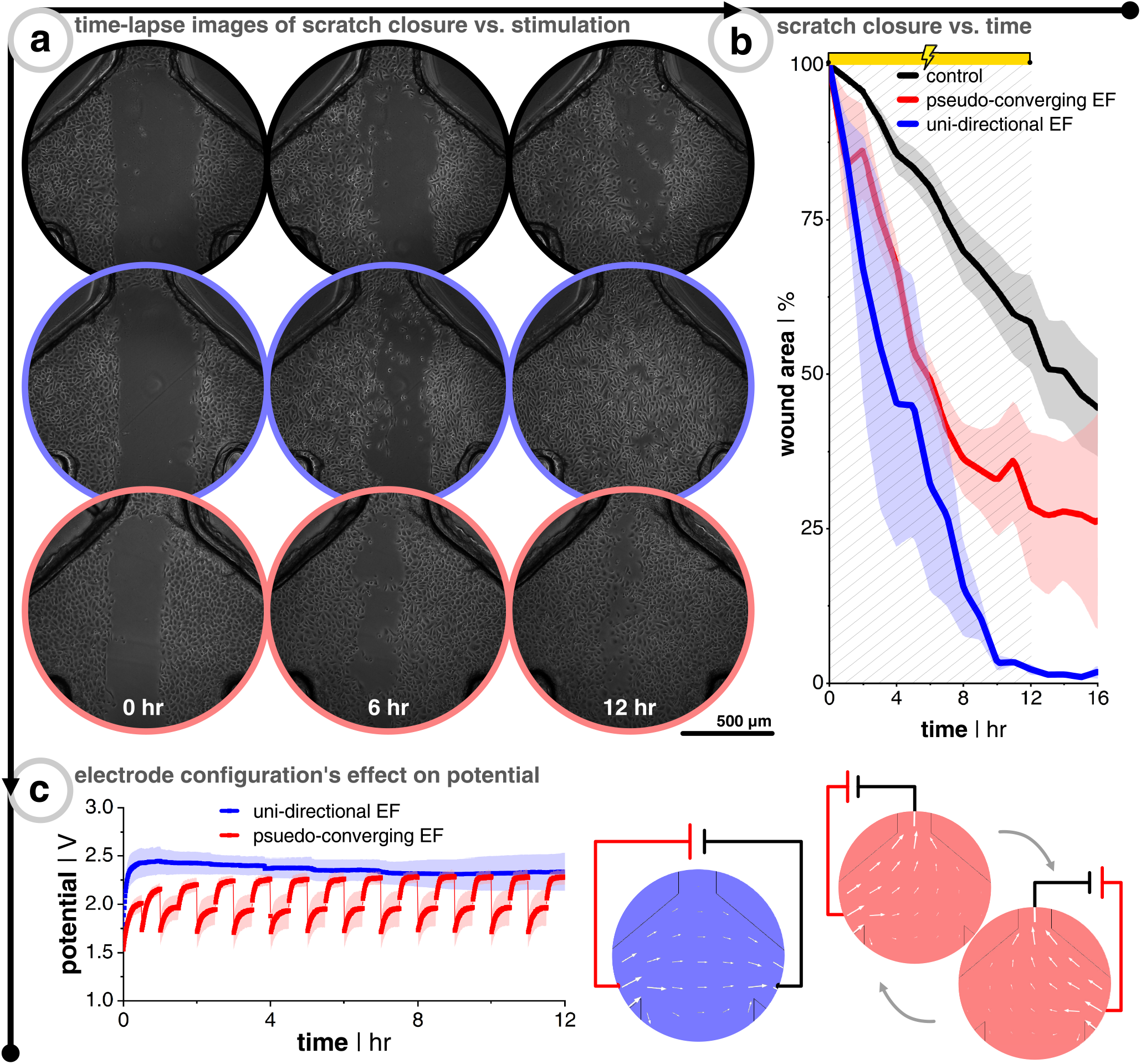
Electrotaxis scratch assay of healthy keratinocytes demonstrate faster wound closure with stimulation. (a) Time-lapse images during 12 hr DC stimulation for non-stimulated control (black, upper panel), uni-directional EF (blue, middle panel), and pseudo-converging EF (red, lower panel). (b) Wound closure over time where the wound area is normalized to the first image (n = 3 for all conditions). (c) example profiles of potential versus time for both electrode configurations. Note that for the converging case that an extra anode is connected (compared to uni-directional case) and a relay switches the anode every 30 min to push cells from both sides of the scratch, as well as passively recharge the unconnected anode with ions from the media.

### DC stimulation promotes recovery of inhibited keratinocyte wound closure motility

In order to explore the hypothesis that EF stimulation not only accelerates scratch closure for healthy cells, but also is of relevance for patients with impaired wound healing, it is compelling to establish a protocol to mimic the less motile wound closure phenotype of diabetic wounds. Once established, this could be translated to testing inhibited cells under direct current EF to see if it helps recover the lost motility. Two approaches to model diabetes are tackled in this paper. The first is to subject the seeded keratinocytes to a hyperglycemic environment (i.e., high glucose concentration), which restrains migration speed via sequential suppression of the p-Stat-1 pathway and the *α*2*β* 1-integrin-mediated MMP-1 pathway.42 Another approach is to inhibit the p38 mitogen-activated protein kinase (MAPK) pathway. This pathway is directly involved in the transition of keratinocytes from cells who are destined to terminally differentiate into the outermost skin layer (i.e., stratum corneum), and help transform them into highly migratory cells upon wounding, which is part of the regeneratory process.^43, 44^ For diabetic wounds, the p38/MAPK pathway has shown to be down-regulated in high glucose environments, which would prevent this transition. Thus, if this pathway can be restored (up-regulated), then it might be possible to restore the cell migration via an autophagy-dependent manner.^45^ Jiang *et al*. showed that down-regulation of CD9, a gene encoding protein involved in cell motility, promotes keratinocyte migration. Also, p38/MAPK inhibition increases CD9 expression, thus supressing migration.46 Building off that knowledge, it was also shown that keratinocytes subjected to direct current EFs (200mVmm^−1^) will have their CD9 expression down-regulated via the 5′ adenosine monophosphate-activated protein kinase (AMPK) pathway.^47^ These cumulative factors lead us to believe that if keratinocytes are slowed via an inhibited p38/MAPK pathway (increased CD9 expression), then direct current EFs could down-regulate the CD9 and override the migration inhibition via the alternative AMPK pathway.

After seeding keratinocytes so that they were fully confluent the next day, they were either subjected to keratinocyte growth medium spiked with different concentrations of D(+)-glucose (6 to 100 mM), medium spiked with p38/MAPK inhibitor (0.5 to 50 uM), or a combination of both (Fig. 4). The cells were kept in the glucose environment overnight before scratching and imaging, while the p38/MAPK inhibited cells were subjected for 3 hr. Importantly, all inhibitor treatments tested did not affect cell viability even after 24 hr of treatment (Supplemental Fig. 6). Both treatments on their own were successful at slowing down scratch closure, starting at 100 mM for glucose and 25 uM for p38/MAPK inhibitor. The wound closure rate was more than halved compared to the untreated controls (Fig. 4b, black trace, Supplemental Video 3). The combinations of both treatments (the amalgam of 50 or 100 mM glucose and 25 or 50 uM inhibitor) were also successful at reducing migration speed, but were much less consistent amongst replicates compared to the single treatments (Supplemental Video 4). All things considered, it was decided that the effectiveness and reproducibility of p38/MAPK inhibitor (25 uM) was the best condition moving forward since it decouples possible compounding issues of dual treatments, in addition to targeting a specific pathway that is potentially more directly linked to electrotaxis.

**Figure 4.**
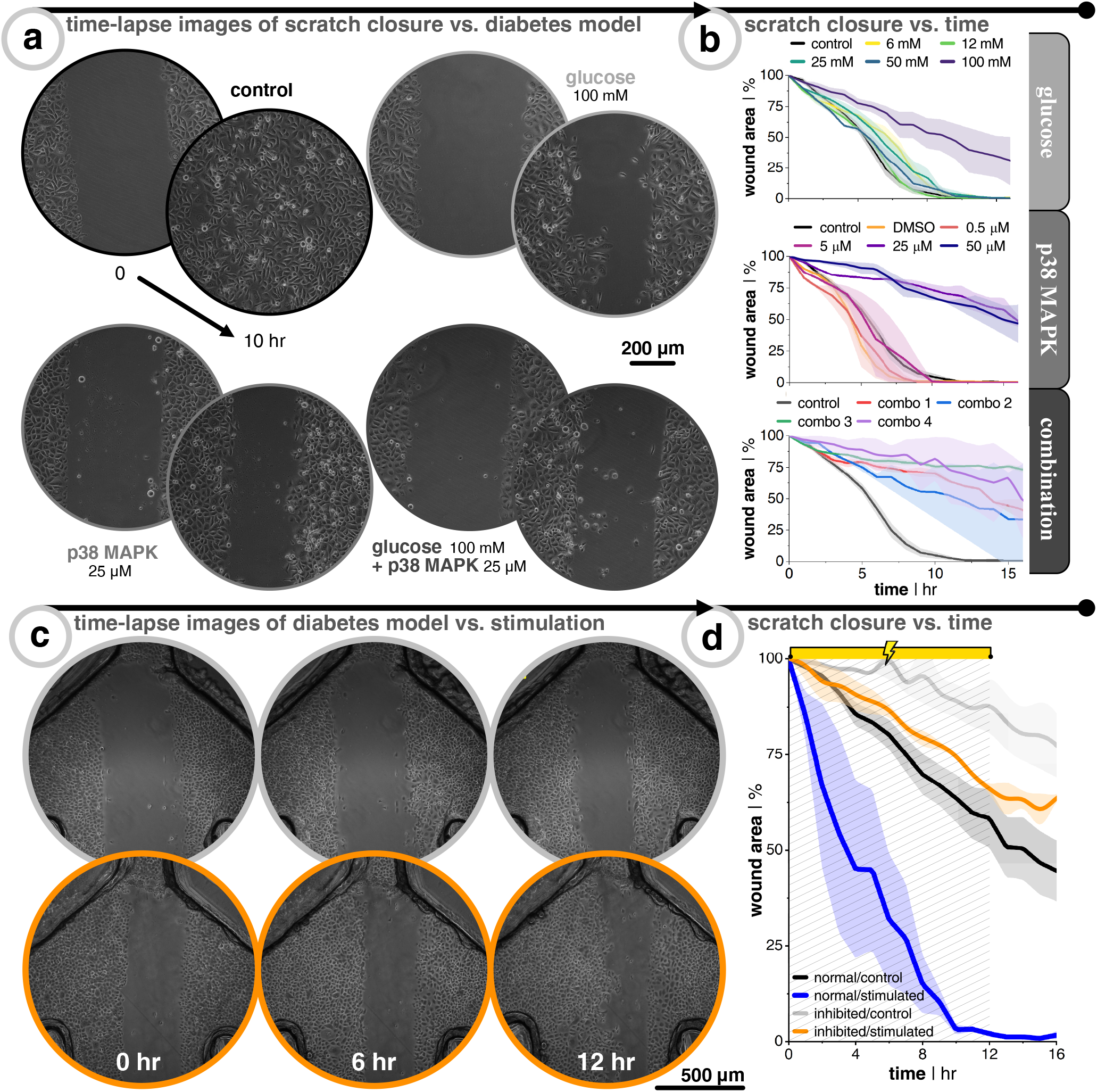
Inhibitory treatments of keratinocytes slow down migration and electrical stimulation helps recover lost motility. (a) Keratinocytes seeded on a 12-well plate are subjected to normal growth media (control), media with D(+)-glucose, media with p38/MAPK inhibitor, or a combination of both. After treatment, a scratch assay is performed and time-lapse images are taken. (b) Wound closure over time where the wound area is normalized to the first image (n = 3 for all conditions). Note that p38/MAPK inhibitor stock solution is diluted with dimethyl sulfoxide (DMSO). Therefore, a control group was added with the same final concentration of DMSO that was used in all inhibitor cases in order to account for DMSO’s effect on scratch closure. (c) Time-lapse images during 12 hour DC stimulation for non-stimulated control (grey) and uni-directional EF (orange). (d) Wound closure over time where the wound area is normalized to the first image (n = 3 for all conditions). Blue and black traces are a carryover from Fig. 3 in order to facilitate comparison with healthy keratinocytes.

Importantly, the positive effect of EF stimulation on wound closure was demonstrated here to hold true also for p38/MAPK-inhibited keratinocytes. It was clear that direct current EF guidance cues help close the wound faster than non-stimulated controls (Fig. 4). After 12 hr of uni-directional EF stimulation, inhibited cells (n = 3, in orange) was ≈ 34% closed compared to only ≈ 12% closed for non-stimulated controls (n = 3, in grey). To put this ~ 3X increase in closure speed into perspective, it helped the inhibited cells to nearly recover to the same wound closure speed as once healthy keratinocytes (Fig. 4b, orange vs. black traces, Supplemental Video 5). Our findings support the idea that electrical field guidance can act to support faster wound closure and, in particular, the potential relevance for addressing impaired wound healing associated with diabetes.

## Discussion

The harmonization of microfluidic design, electrode material choice, and assembly protocol that were shown in this work provides a new platform for exploring the effects of electric fields on wounded cells of choice. The platform is based on readily-available materials and commonly-used prototyping equipment, and could easily be manufactured and customized by others according to our protocols. Below we discuss three major topics that were leveraged in this work, their key parameters, and the future outlook: DC electrode materials, guidance cues, and disease models.

A major hurdled obstacle that we report here is the application of electrodes that do not require salt bridges, which allows for easier translation to 3D models.^28^ The use of non-metal, supercapacitive hydrogel electrodes facilitates not only a compact design in this 2D assay work, but ease of translation to more 3D architectures, and importantly, are an enabling technology for translation into a clinically useful device. Although, it is of critical importance to focus on the electrode’s charge storage limitations and the stimulation protocol’s total charge to be delivered. The pseudocapacitor hydrogel electrode used here has a relatively high charge storage capacity (CSC ≃ 40 to 50 mCcm^−2^).^31^ This CSC is crucial for determining over how long the current can be delivered capacitively (*t* = *q*/*i*, where *q* is charge stored in electrode and *i* is the input current) before it shifts to a mostly faradaic current, which is reliant on pH-shifting electrochemical reactions. This is why it is imperative to also account for the amount of pH buffering agent (molarity). On the one hand, the availability of nearby buffering agents in *ex vivo* or *in vivo* constructs might not be as abundant as in carefully engineered *in vitro* systems, however on the other hand, perfusion in functional tissue can help supply reinforcing buffering agents. Additionally, we demonstrate that over-polarized electrodes after stimulation are able to recharge with ions from the surrounding biological electrolyte and that switching between two anodes mitigated faradaic-induced pH shifts (Supplemental Fig. 3e). This opens up the possibilities to employ multi-electrode arrayed anodes and cathodes that have sub-groups actively discharge (i.e., unipolar, constant current), while other sub-groups passively recharge (i.e., disconnected), and then cyclicly interchange between these two sub-groups. It should here be noted that metal electrodes typically do not possess this ability to recharge, as their DC charge injection involves corrosion, a reaction that cannot easily be reversed. In addition, corrosion typically elute toxic metal ions into the tissue. Meanwhile, the metal-free electrodes used here work with ions available in abundance in the biological electrolyte.

Right now, the fields of bioelectronics and wound healing are just ‘scratching’ the surface of how best to use guidance cues to control cell collectives. This work unveiled the surprising result that constantly pushing a wound to close from one side via EF guidance cues was more effective than alternatingly directing the wound to close from both sides. This notion can be leveraged when designing future electrical wound dressings since a truly converging EF design would require the problematic task of placing a cathode within the wound’s exudate. Furthermore, with an extensive variety of wound morphologies, the centrally-place cathode becomes more difficult to standardize, whereas this burden would be less so with the simplicity of fixing an electrode on opposite sides of the wound. Here, additional electrode montages could be employed to focus on wound closure from different sides or edges. With that being said, there is still a need to further dive into how guidance cues impact group migration. Recently, Shim *et al*. demonstrated that increasing levels of calcium proportionally increases cadherin-mediated cell-cell adhesion strength in between adjacent keratinocytes, reduces average cell migration speed, and dampens the effect external direct current EF has on directionality, which goes to show there is an interplay of cell-cell forces, cell-substrate forces, and external guidance cues in effective collective migration control.^48^ In conjunction with bioelectric cues, a mixture of mechanical cues (e.g., extracellular matrix coatings) and chemical cues (e.g., passive or active flowing of pro- or anti-migratory soluble factors) are compelling to investigate and straight-forward to implement in a platform such as the one presented here. Also, it is feasible to leverage microfluidic laminar flow to cleverly focus flow^49^ of compounds of interest in a spatial manner (also possible here in Supplemental Fig. 2). These external cue-mediated 2D sheet migration insights may unveil new mechanisms, and are of importance for translating findings from single cells to more complex and tissue-relevant architectures.

Modeling a disease *in vitro* has its limitations, but offers opportunities to study individualistic cause and effects. Diabetes *in vitro* models have been thoroughly explored in the aspects of neuropathy^50^, pregnancy^51^, and wound healing^45^. Hyperglycemic *in vitro* studies of wound healing rates seem to vary dependent on epidermal growth factor (EGF) concentration in glucose-spiked media, which suggests that solely using glucose concentration as the independent variable is not targeted enough.^52^ However, targeting downstream effected pathways of hyperglycemia, like the p38/MAPK pathway, offers a more directed approach to induce a diabetic-like state in the most wound-prevalent cells, keratinocytes. Prior studies have generated evidence that other pathways also play an influential role in diabetic wounds, including the diacylglycerol pathway, hexosamine pathway, protein kinase C pathway, and polyol pathway.^53^ This illustrates that there is more room to explore the impact of multiple guidance cues on a variety of relevant diabetic wound pathways at the foundational level. More ambiguous diseases, like systemic lupus erythematosus (SLE), also have affected wound healing. Diseases like SLE stand to benefit from more fundamental research on responsible pathways that could potentially be overridden by electrical stimulation in order to improve their typically poor healing of wounds. There is also evidence that transcutaneous electrical stimulation promotes healing of intact skin for such ailments like pressure ulcers of paraplegic individuals.^54^ However, for all these aforementioned diseases there needs to be more exploration into the mechanistic effects of electrical stimulation and more encompassing dose-response investigations from the collective cell group scale all the way up to the organ level.

The platform developed here allowed us to lay the foundation for a future wound healing concept based on electrical stimulation from supercapacitive non-metal electrodes. We demonstrate the working principle of this concept using culture models of skin wounds, and show that EF guidance cues can increase the wound closure speed up to 3X, in comparison to unstimulated controls. We furthermore show that effective wound closing stimulation relies on a carefully controlled environment, dosage, and directionality of the electric field. Under the condition that these factors can also be accounted for in real skin wounds, we are convinced that electrical stimulation could contribute as an additional guidance cue in regenerating tissue and thereby promote faster re-epithelialization.

## Methods

All chemicals were purchased from Sigma Aldrich, unless otherwise noted.

### Finite element analysis of electric field distribution and joule heating

COMSOL Multiphysics^®^ software (version 5.3) was used to simulate both EF distribution and joule heating, using the Electric Currents and Heat Transfer modules, respectively. SolidWorks (version 2021) was used to design the three-dimensional models used for COMSOL. For EF distribution, electrodes sat on top of the reservoirs and were modeled to have electrical conductivity of PEDOT:PSS hydrogels (σ = 2000 Sm^−1^).^55^ The media was modeled after 1X (i.e., 10mM) phosphate-buffered saline (PBS, σ = 1.54 Sm^−1^). Only the form factor of the media and the electrodes were modeled. The input current density (placed at face of anode(s)) was sweeped in order to identify at which current the desired EF strength would be reached at the center of the channel. The cathode was set to a potential of 0 V. For joule heating, the same electrical conditions were implemented, but changed to a time-dependent solver. This model also included modeling full media-filled reservoirs and the plastic substrate and lid. The following values were used for heat transfer-related material properties (values are from COMSOL’s material database unless otherwise cited): PBS (k = 2Wm^−1^ K^−1^, *ρ* = 1000 kg m^−3^, *C_p_* = 4 J kg^−1^ K^−1^),^56^ acrylic (k = 0.19 W m^−1^ K^−1^, *ρ* = 1190 kg m^−3^, *C_p_* = 1420 J kg^−1^ K^−1^, σ = 1 × 10^−14^ S m^−1^) PEDOT:PSS (k = 0.348 W m^−1^K^−1^,^57^ *ρ* = 1060 kg m^−3^,^58^ *C_p_* = 1415 J kg^−1^ K^−1^).^59^ The initial conditions include a convective heat flux of external temperature of 310.15 K with heat transfer coefficient of 5 W m^−2^ K^−1^, diffusive surface of all plastic components with surface emissivity of 0.95 and ambient temperature of 310.15 K. The fluid was modeled to have zero velocity and a pressure of 1 atm.

### Microfluidic fabrication

All components of the microfluidic device, besides the sterile polystyrene dish, are fabricated with a 30 W carbon dioxide (CO_2_) laser (Universal Laser Systems, VLS 2.30). Specifically, 7.5 W at 70mm/s was used for a kiss-cut and 24 W at 70mm/s was used for a through all-cut for the acrylic-based double-sided pressure-sensitive adhesive (Adhesives Research, 90445Q). The bottom-side liner (i.e., without the kiss-cut) is peeled off to expose the bottom-side adhesive, then it is pressure bonded by hand to a new Petri dish. Batches of dish/adhesive were placed in a vacuum desiccator overnight to remove any air bubbles that occurred during bonding. These are stored on the shelf until further use. The acrylic (Modulor, Germany) two-part lid included a thin 0.5mm base that only has fluidic vias and a thicker 8.0mm reservoir-defining layer. These two parts were solvent bonded together using dichloromethane (Modulor, Germany).

### Electrode fabrication

The fabrication of laser-induced graphene (LIG) electrodes coated with pure PEDOT:PSS hydrogels was recently established by our group.^31^ In short, a (CO_2_) laser (same as above) is used to carbonize a polyimide sheet (Kapton HN, 75 μm thick) with a rasterization protocol at 4.8 W and 15.2 mm/s. The freshly carbonized LIG was air plasma-treated (Femto, Diener Electronics) for 5 min at 100 W and 10 sccm to make it more hydrophilic and functionalizable. They were then soaked in 10%w/v hexamethylenediamine (HMDA) for 4 hr at room temperature. After washing with DI water and drying with nitrogen, they were dip-coated (Nadetech ND-DC Dip Coater) in a 1 %w/v hydrophilic polyurethane solution in 90 % ethanol (AdvanSource, HydroMed D3) with a retraction speed of 100 mm min^−1^, then subsequently placed on a hot plate for 1 hr. Electrode connection lines (i.e., between electrical bump pad and electroactive area) were coated with a acrylate-based varnish (Essence 2 in 1, Cosnova). PEDOT:PSS dispersion (1.3 %) with 15 % dimethyl sulfoxide was drop-casted (200 μL onto the 12 mm diameter LIG electrodes after being placed onto a hot plate (60 °C overnight, then 130 °C for 90 min). Electrodes were stored in 1X PBS (Sigma Aldrich, P3813). Electrical connections were done with a pogo-pin assembly (Mill-Max, 858-22-002-30-001101) fitted with M3 threaded inserts. Custom polyethylene terephthalate glycol (PET-G) 3D-printed (Prusa Research, i3 MK3S) parts were made for compression clamping of the pogo pin assembly (seen in green in Fig. 1) and for consistent angled alignment of electrodes into reservoirs (seen in orange in Fig. 1).

### Coating of pH-sensitive polymer

The polyaniline (PANI) is polymerized via chemical oxidation^60^ by mixing equal volume and molarity of pre-chilled aniline monomer (1.42M) in hydrochloric acid (HCl, (1.42M)) with pre-chilled ammonium persulfate (APS, (1.42M)) in deionized water and depositing this polymerizing mixture onto the open microchannels (Supplemental Video 1). Note that the aniline solution and the APS solution are prepared fresh and are stored at −20 °C for 30 min before mixing to slow down the polymerization in order to facilitate sufficient time for deposition. The mixture turns from a colorless solution into a deep black hydrogel over the next 30 min (Supplemental Video 1). The *in situ* polymerization embeds itself into the Petri dish polystyrene such that unbound PANI hydrogel can be washed away leaving behind a thin layer of green PANI (i.e., emeraldine salt).

For calibration, PANI was coated onto small polystyrene Petri dishes (35 mm diameter, Falcon TC-treated). Stock pH buffers ranging from pH 2.69 to 11.25 were made using different combinations of spiking hydrochloric acid (HCl) or sodium hydroxide (NaOH) into primary salt solutions made of either potassium hydrogen phthalate (KHP), potassium dihydrogen phosphate (KH_2_PO_4_), sodium tetraborate (Na_2_B_4_O_7_), or sodium bicarbonate (NaHCO_3_). A benchtop pH meter was used to verify all pH values (VWR pHenomenal pH 1100L). After depositing 1 mL of the pH buffer into seperate PANI-coated dishes, they were imaged in an incubated inverted microscope (Zeiss Axio Observer) with a 20X objective at 37 °C. Images were processed through a Python script to report a hue value. Specifically, the script averages RGB values across the image and reports a corresponding hue. Based on the maximum and minimum values of RGB, the hue is then calculated by using the Python module “colorsys” to change from RGB to HSV coordinates. These hue values were correlated to the known pH values to generate a calibration curve. The curve was fitted with a bi-dose response sigmoidal curve (Origin 2021).

For coating PANI in the DC stimulation device, the polymerization and casting process is the same, but now onto the polystyrene-exposed parts of the microfluidic-defining adhesive (Supplemental Video 1). After washing away unbound PANI, the liner was then removed and lid was subsequently added, just like in Fig. 1e, steps 5 & 6. PBS (1X) was degassed in a vacuum desiccator for 30 min to minimize bubble formation in the microchannel. Then, degassed PBS and electrodes were added the device. Imaging was performed in an incubated microscope over the course of 20-plus straight hours of monophasic DC stimulation (25 μA using a potentiostat/galvanostat (Metrohm, Autolab PGSTAT204). For the pseudo-converging EF case, a custom-built relay was used between the current source’s working electrode lead and the two anodes.

### Culturing keratinocytes

Human epidermal keratinocytes immortalized with HPV-16 E6/E7 were courtesy of Prof. Dr. rer. nat. Thorsten Steinberg (Department of Dental, Oral and Jaw Medicine; University Clinics of Freiburg). Keratinocytes were cultured throughout experiments in serum-free keratinocytes growth medium (KGM2, PromoCell, #C-39016) supplemented with cocktail of factors and CaCl_2_ provided by the same manufacturer (SupplementMix, PromoCell, #C-20011), as well as neomycin (Sigma-Aldrich, #N1142) at final concentration 20 μgmL^−1^ and kanamycin (Sigma-Aldrich, #K0254) at final concentration 100 μgmL^−1^. Cell culture was incubated at 37 °C and 5% CO_2_ at 95% humidity and routinely passaged when 80 to 90 % confluency was reached. Growth medium was exchanged three times per week. For the experiments, keratinocytes were used between passages 34 to 49.

### Treatments of keratinocytes to mimic diabetic phenotype

For experiments with elevated glucose concentrations, a 1M aqueous stock solution of D(+)-glucose (Sigma Aldrich, #G7021) was added to the culture medium to achieve a desired concentration (6, 12, 25, 50, or 100mM). The glucose-rich medium was prepared fresh each time before treatment. Confluent cell layers were treated for 24 hr before proceeding to the wounding.^42^

For experiments mimicking diabetic wound environment, a 25 mM stock solution of p38-MAPK inhibitor (Cell Signalling via Selleck Chem, Adezmapimod - SB203580, #S1076) in DMSO was added to the culture medium to achieve a desired concentration (0.5, 5, 25, or 50μM). The inhibitor-containing medium was prepared fresh the same day as treatment. A control condition to test the effect of DMSO on cell viability and migration was done by using the same final DMSO concentration (0.1 %v/v) as above, but without inhibitor. Confluent cell layers were treated for 3 hr before proceeding to the wounding.^45^

In order to assess viability of cells after treatment, live/dead cell double staining was performed with SYTO 16 (Invitrogen, #S7578) and propidium iodide (Invitrogen, #P1304MP). For staining, culture medium containing both dyes with a final concentration of 500nM each was prepared. Cells were protected from light and incubated at 37 °C for 30 min, then washed with 37 °C PBS and imaged with incubated inverted microscope (Zeiss Axio Observer).

### Seeding cells onto devices, wounding monolayer, and device assembly

Before seeding onto scratch assay devices, the devices were air plasma-treated (30 W, 3 min, 10 sccm) to improve cell adhesion to the substrate. To prepare for scratch assays, keratinocytes were detached from culture flasks by incubation with 0.05% trypsin and 0.02 % EDTA solution (Sigma Alrich, #T3924) at 37 °C for 5 min. To neutralize trypsin, medium containing 10% fetal bovine serum (Sigma Aldrich, #F0804) was used. Harvested cells were centrifuged (1200RPM for 10 min) and resuspended at 4.5 × 10^6^ cells mL^−1^. Cell suspensions (100 μL) were spotted directly over open microchannels (see Fig. 1b, step 3) and incubated for 3 hr to allow for cell attachment. Afterwards, the excess of cells was washed away with 37 °C PBS solution and aspirated before adding 10 mL of fresh growth medium. Cell seeding concentration was titrated in order to find the optimal amount so that devices were fully confluent the next day (Supplmental Fig. 5).

Monolayers were scratched using a sterile p10 pipette tip (≃ ∅ 700 um) that was connected to a vacuum aspirator (Vacusip, Integra Biosciences). The motion of the scratch was always done starting at the base of the microchannel scratch alley and finished where the four channels converge. After the scratch, the media was aspirated, then devices were washed with sterile PBS (1X), then fresh growth media was added, and finally placed back in the incubator for 4 hr. In the meantime, the acrylic lid, electrodes, 3D printed adapters and wires are washed with 70% ethanol and then further sterilized in the S1 cell culture hood (Safe 2020, Thermo Scientific) via UV-treatment for 1 hr. After incubation, media was aspirated until only a small amount of media resided in the microchannels and leaving the liner as dry as possible in order to minimize the probability of the fluid transferring onto the dry adhesive. The liner was then peeled, the two-part acrylic lid was aligned and fixed using alignment marks, and media immediately replenished by initially flowing 100 μL directly into the microchannels to displace trapped air. The rest of the reservoirs were filled using a standard serological pipette. The electrodes were assembled and placed into the reservoir, and the corresponding wires were routed through the lid, which was applied to prevent evaporation.

### Imaging and direct current stimulation

Seeded and assembled devices were placed in an incubated inverted microscope (Zeiss Axio Observer with Definite Focus 2) and maintained at 37 °C and 5% CO_2_. Phase-contrast images were acquired every 10 min using a 5X objective in order to capture the entire microfluidic network. The DC stimulation was carried out the exact same way as described in the preliminary pH-monitoring experiments (e.g., constant current using potentiostat/galvanostat - Autolab PGSTAT204). Output images were put through an ImageJ plugin in order to quantify the scratch area closure over time.^61^

## Acknowledgements

This project was funded by the European Research Council (ERC) under the European Union’s Horizon 2020 Research and Innovation program under grant agreement (No. 759655, SPEEDER). This work furthermore received support by BrainLinks-BrainTools, Cluster of Excellence funded by the German Research Foundation (DFG, EXC 1086), currently funded by the Federal Ministry of Economics, Science and Arts of Baden Württemberg within the sustainability program for projects of the excellence initiative. We would also like to acknowledge the Wissenschaftlige Gesellschaft Freiburg for added financial support and the Deutscher Akademischer Austauschdienst (specifically, the International Association for the Exchange of Students for Technical Experience - IAESTE) for facilitating financial support for the author A.K. to work in our group on summer exchange. Finally, we would like to thank Dr. Christian Boehler for his fruitful advice on pH-monitoring experiments and Mrs. Ute Riede for her immeasurable help with supporting cell culture maintenance and seeding protocols.

## Author contributions statement

S.S., M.A., and A.S. conceived the experiments, S.S., A.S., A.K., and N.J. conducted the experiments, S.S., A.K., L.M., J.L., and M.A. analyzed the results, S.S. made the figures. All authors reviewed the manuscript.

