## Supplemental Figures for "Bioelectronic microfluidic wound healing"

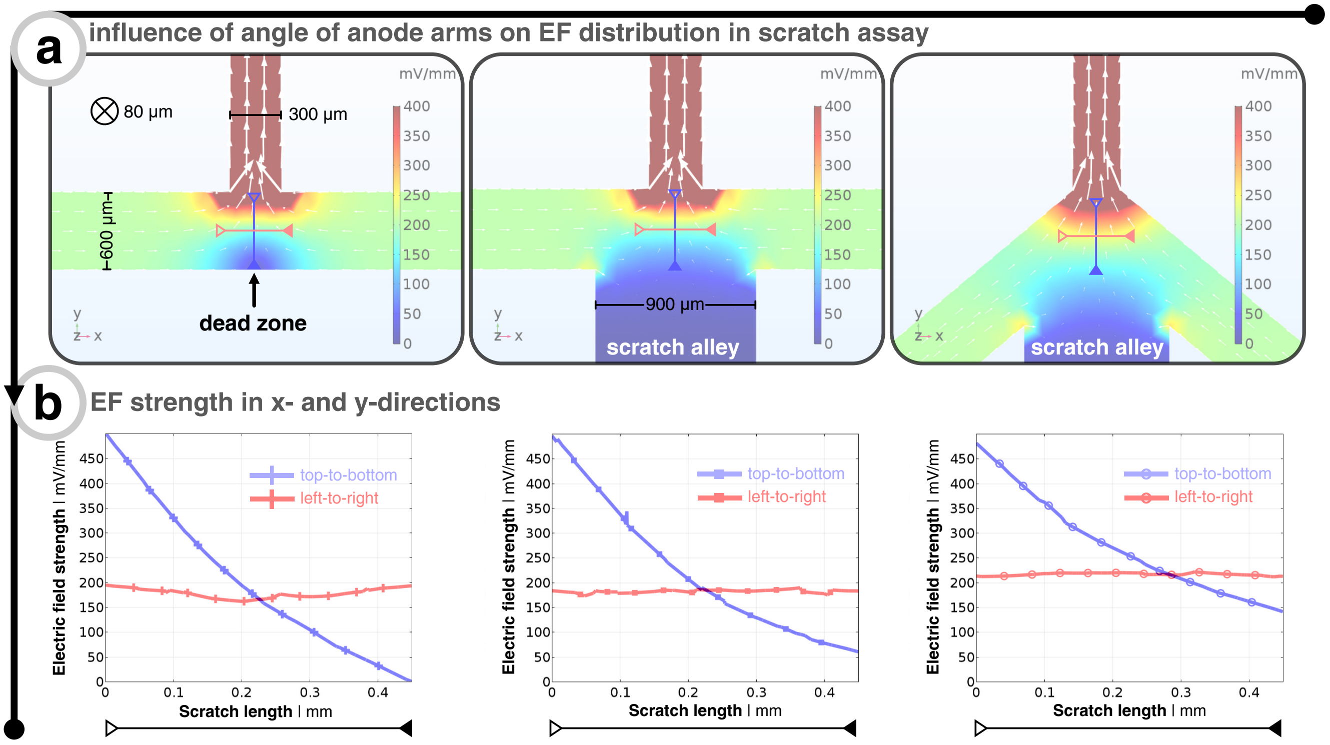


**Figure 1. Finite element analysis reveals the effect of the angle of converging anode arms and scratch alley on EF distribution**. (a) Three different configurations of the angle of the left and right microfluidic branches in relation to the top cathode branch and the existence of a scratch alley or not. Note that for all cases, the channel dimensions are consistent between models, there are two shorted anodes (one on both sides of the converging zone), and a single cathode connected to the top channel. (b) Electric field strength of blue and red line sections from (a). The blue line starts from entry of top channel and points downward. The red line starts left of the theoretical scratch zone and ends on the right of said zone. Forcing converging current into the top channel of a T-junction creates a dead zone, seen in left column and blue trace. Adding a scratch alley (middle column) helps minimize the dead zone effect seen in blue trace. The angled anode branches helps even further.

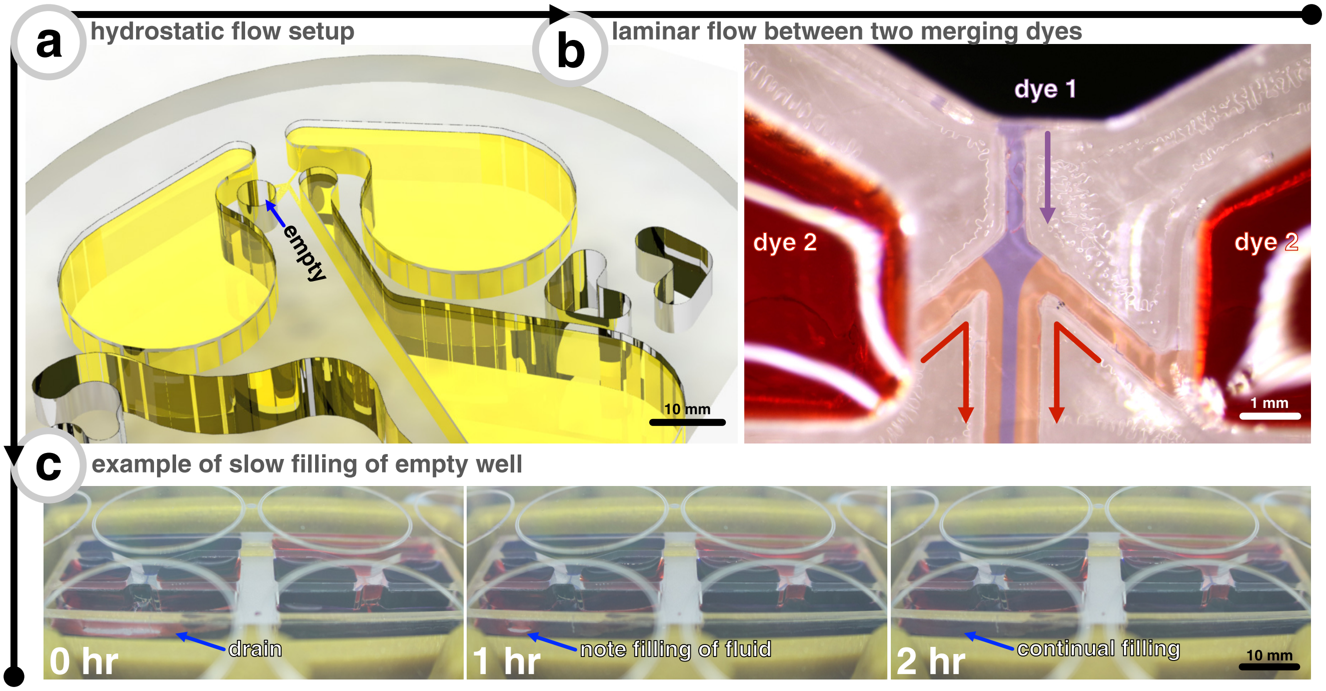


**Figure 2. Hydrostatic flow option in microfluidic scratch assay device**. (a) setup of how hydrostatic pressure can be used for passive perfusive flow where the three electrode wells are full and the fourth drain well is empty. (b) adding dye 1 (Brilliant Black BN) in top well and dye 2 (Congo Red) into left and right wells show how flow is laminar when fluids used have different properties. (c) time-lapse example of how fourth drain well slowly fills due to hydrostatic flow.

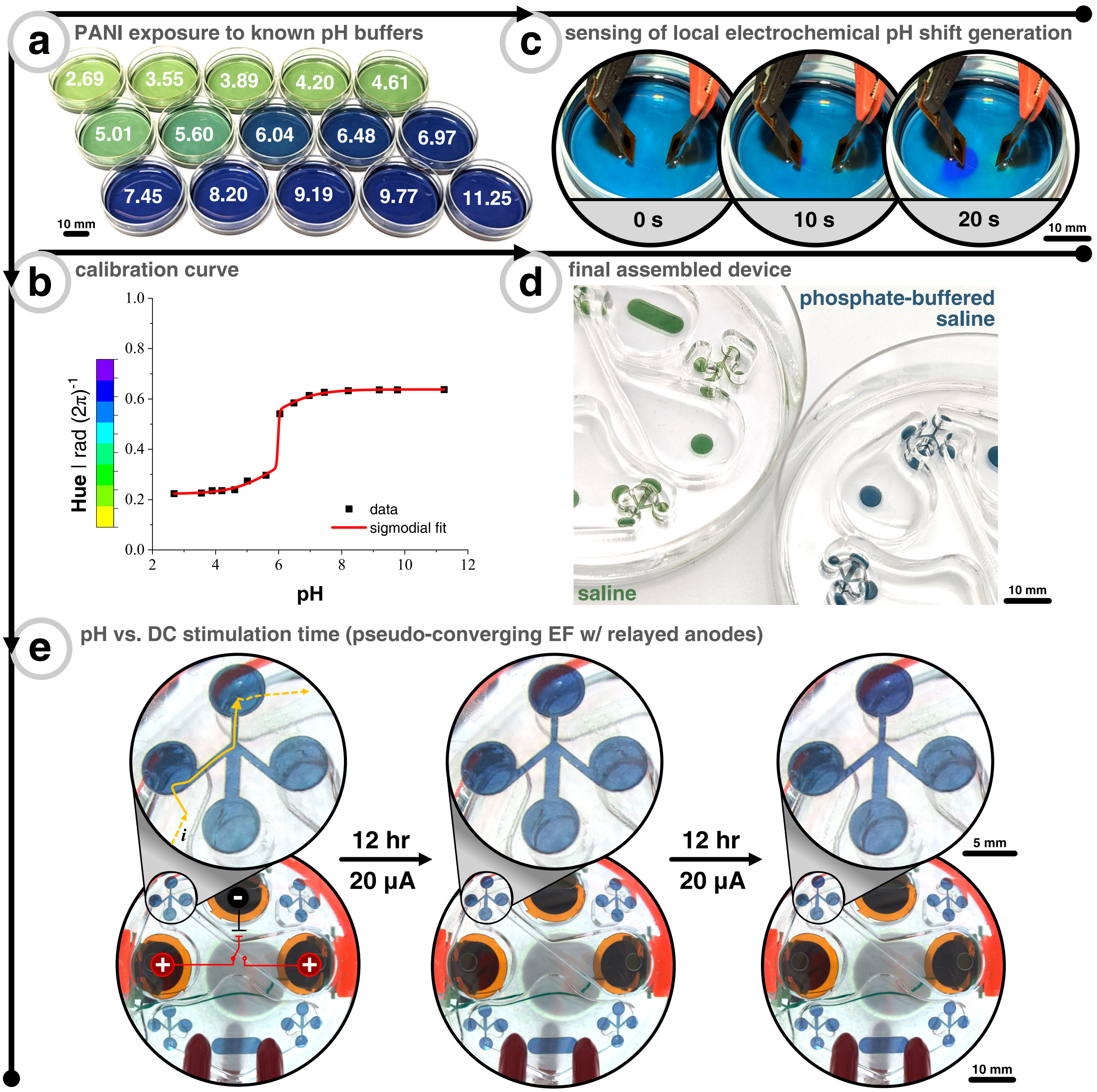


**Figure 3. Calibration curve of PANI color versus known pH buffers**. (a) PANI-coated Petri dishes (35 mm diameter) subjected to 15 different known pH buffers that were previously measured with a benchtop pH meter. (b) Calibration curve when hue of PANI color changes when subjected to wide range of pH buffers. Note that calibration images were taken in the same manner as they were for Fig. 2 (main text). In short, in an incubated transmission microscope with 5X objective with same lamp and exposure settings. The curve fitting was with a "bi-dose response" sigmoidal fit. (c) Demonstration of how the pH-sensitivity of PANI can be used to visualize pH changes induced by DC stimulation. Cathode is on the left and anode on the right. Electrodes used are the same materials, but smaller (3 mm diameter), as the electrodes used throughout the paper. Solution is unbuffered 0.9% NaCl and current is constant at 0.40 mA. (d) Example of final assembly of PANI-coated scratch assay devices when filled with saline (pH = 5.4) and phosphate-buffered saline (pH = 7.4). (e) Time-lapse images of PANI-coated scratch assay device with pseudo-converging EF. Note that stimulation protocol is identical to that of Fig. 3c (main text). The relayed nature of switching between left and right anodes prevent extended time in faradaic current regime, which allows for minimal anode pH shifts (i.e., no noticeable lowering of pH at anodes) for the entire 24 hr of DC stimulation. However, since the cathode is not switched, the cathode well gets more basic (i.e., purple).

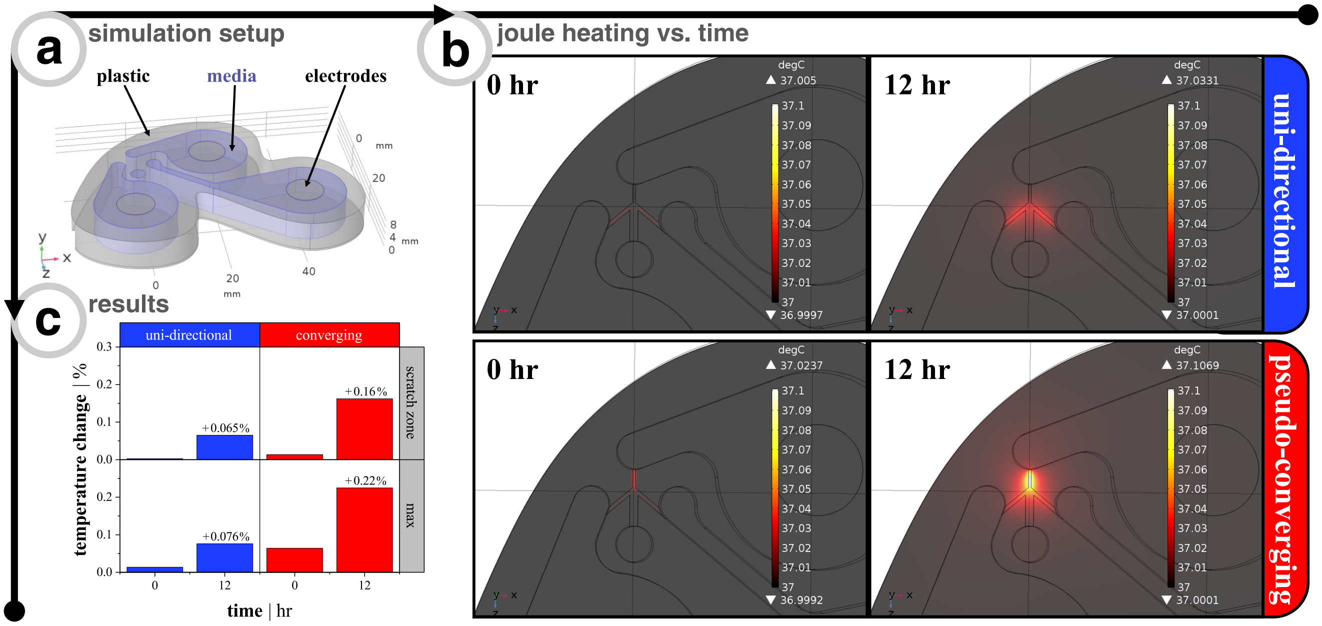


**Figure 4.** **Simulations of joule heating due to direct current (DC) stimulation**. (a) Setup of simulation where the blue shows the electrolyte filled reservoirs surrounded by plastic (i.e., acrylic) and the electrodes sitting on top of the electrolyte. Microchannels that connect the reservoirs are not known here, but can be inferred from Fig. 2 and Fig. 3 (both from the main text). The surrounding temperature boundary condition was set to 37 °C, which serves as the reference point for percent change of temperature. (b) Snapshots of joule heating expected in microchannel for both electrode configurations after 12 hr of constant current stimulation. (c) Simulation results showing minimal joule heating of electrolyte even after long-term stimulation. Whether in the small channels or scratch zone, the heating generated is within 1/10 of a degree Celsius, which is negligible.

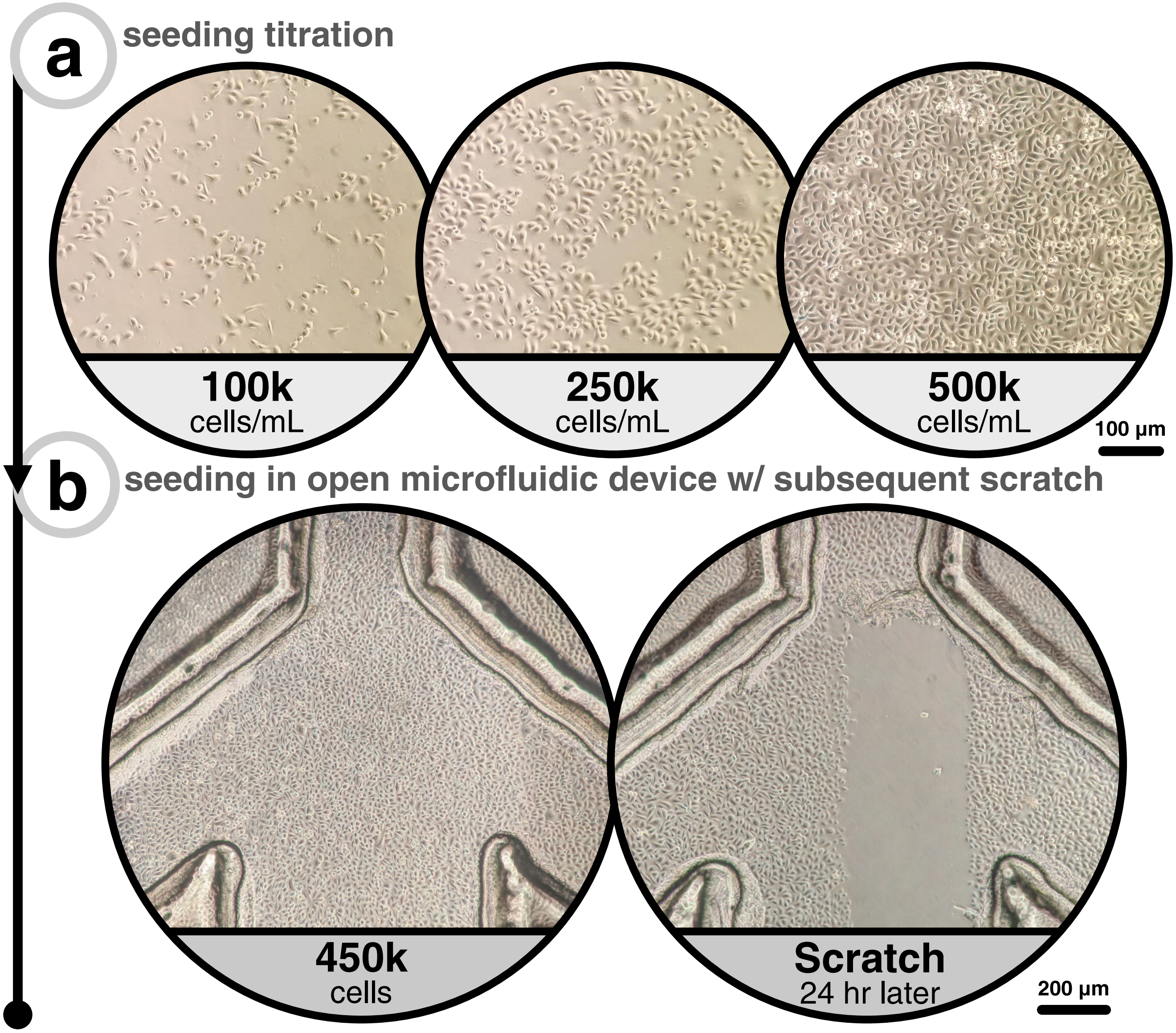


**Figure 5. Seeding titration of keratinocytes to find full confluency**. (a) Seeding titration of cell suspension density to find out spatial adherence density 3 hr after spotting cell suspension on substrate. Reminder that the goal was to have a fully confluent layer of cells 24 hr later for scratch experiments. (b) Final cell suspension density ($4.5 \times 10^6$ cells $mL^{-1}$ in 100 uL) selected for all scratch assay device experiments, and after 24 hr this is an example image immediately after a typical scratch.

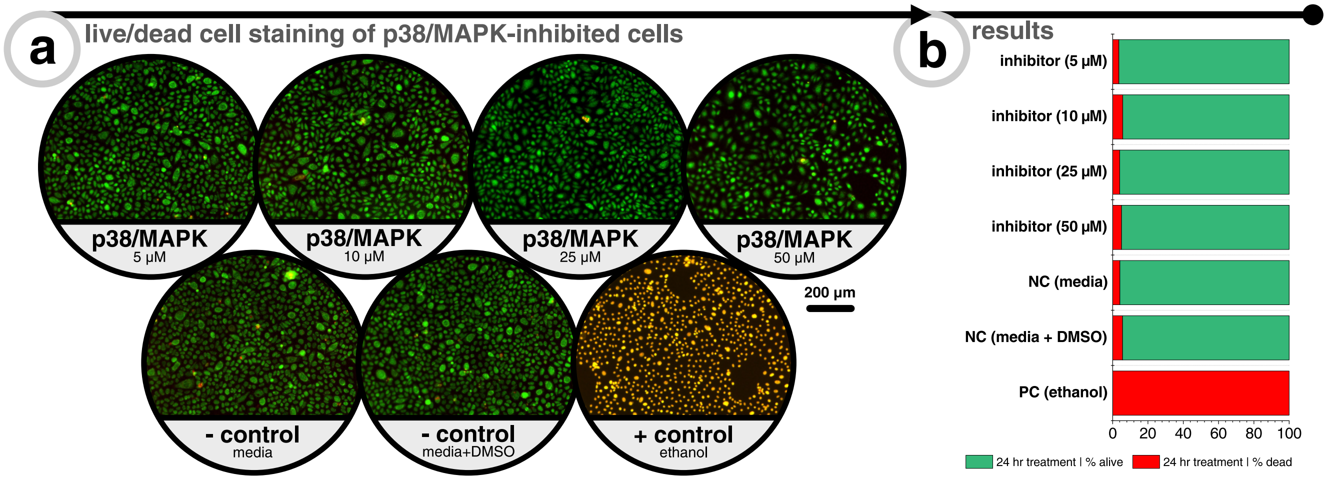


**Figure 6. p38/MAPK-inhibitor treatment does not impact cell viability**. (a) Example images of live/dead (SYTO 16 & propidium iodide) cell staining of keratinocytes after 24 hr of treatment. Negative controls were either media or media plus DMSO (with same concentration as inhibitor samples). Positive control was a treatment with 70% ethanol. (b) Results of live/dead staining where green and red signifies percentage of total population that is live and dead, respectively.
